# Multi-omics analysis of aggregative multicellularity

**DOI:** 10.1101/2024.03.19.585704

**Authors:** Bart Edelbroek, Jakub Orzechowski Westholm, Jonas Bergquist, Fredrik Söderbom

**Affiliations:** Department of Cell and Molecular Biology, BMC, Uppsala University, SE-751 24, Uppsala, Sweden; Department of Biochemistry and Biophysics, National Bioinformatics Infrastructure Sweden, Science for Life Laboratory, Stockholm University, Stockholm, Sweden; Department of Chemistry-BMC, Analytical Chemistry and Neurochemistry, Uppsala University, Uppsala, Sweden

**Author notes:** corresponding author(s): Bart Edelbroek, Fredrik Söderbom.

**Keywords:** Proteomics, Transcriptomics, gene regulation, *Dictyostelium discoideum*, multicellular development, aggregative multicellularity, Amoebozoa, social amoeba

## Abstract

The extent to which mRNA and protein levels or their regulation correlate remains obscure, especially during development. In particular, this is true for organisms that exhibit aggregative multicellularity, such as the social amoeba *Dictyostelium discoideum*. The transcriptome of *D. discoideum* has been thoroughly studied during multicellular development, however the proteome and the correlation to the transcriptome during transition from uni- to multicellular life have not been analyzed in detail. Here, we present the first paired transcriptomics and proteomics developmental time series during aggregative multicellularity. The dataset reveals that mRNA and protein levels correlate highly during growth, but decrease when multicellular development is initiated. This accentuates that transcripts alone cannot accurately describe gene expression. This dataset can therefore be an important resource to study gene expression during aggregative multicellular development, in particular in *D. discoideum*.

## Introduction

One of the pillars of biology is the central dogma, which states that, exceptions aside, the transfer of information from DNA leads through RNA to produce proteins^1^. Messenger RNA (mRNA) is transcribed from DNA and translated into protein in a quantitative manner. It follows that the levels of mRNA and protein are connected to each other. Both are dynamically regulated, and change due to environmental and developmental cues.

Transcriptomic and proteomic analyses have experienced incredible improvement in throughput and accuracy, however sensitivity of proteomics with LC-MS/MS is still lagging behind the sensitivity of transcriptomics by next-generation sequencing^2^. Also, in proteomics the signal cannot be amplified by e.g. PCR. When comparing these two omics approaches, studies based on transcriptomics have increased dramatically relative to studies based on proteomics, reflected by the number of datasets available in different repositories^3,4^

It is not uncommon that mRNA levels are used as a proxy for the levels of the effector molecules – the proteins. There is, however, not always a linear relationship between mRNA and protein, which can be attribute to e.g. differences in translation rates or protein stability^5^. Studying the correlation between mRNA and protein levels is important in order understand to what extent transcriptomics data can be used to predict gene expression^6^.

Organisms across the tree of life have evolved distinct strategies, which control the balance between mRNA and protein^7^. Due to these differences, the mRNA-protein correlation can differ between species, especially those that are phylogenetically distantly related. Additionally, the correlation between mRNA and protein can be different depending on the biological context of the cell. In steady-state cells, at the population level, the mRNA levels and protein levels are expected to be relatively stable, and their correlation high^5^. On the other hand, when the cells are undergoing changes, many genes will be differentially regulated and the correlation might be lower. One example of this is development, where cells undergo major differential gene expression. Here, cells transition to a different behavior or identity through intrinsic or extrinsic signals.

Previously, the relationship between mRNA and protein has been studied by paired transcriptomics and proteomics at specific developmental stages in several eukaryotic organisms^8–11^. One organism with very particular development is the social amoeba *Dictyostelium discoideum*. When the amoebae run out of food, a developmental program is initiated where, *D. discoideum* transitions from free-living to multicellular. First the cells form aggregates of up to 100,000 cells, which then continue to develop into a fruiting body, where dead stalk cells support a ball of spores^12^. This aggregative multicellular development has been studied thoroughly at the RNA level, characterizing the main processes involved in the developmental program, as well as differentiation into specialized cell types at the single-cell level^13–17^. Thus far, the developmental proteome has not been extensively studied, and it remains unknown how well the observed transcriptional changes are reflected at the protein level.

In this study we performed transcriptomics and proteomics analyses at several time points during early development of *D. discoideum*, to elucidate the mRNA and protein levels throughout multicellular aggregation. We confirmed previous findings, which identified differentially regulated genes involved in processes essential for early development. Additionally, we detected many genes that are dynamically regulated, where specific mRNAs can be, for example, upregulated early during development and down regulated at the later stages, and vice versa. However, at the protein level many of the dynamically regulated mRNAs result in linearly regulated proteins. Another observation was that, in general, protein expression is delayed several hours as compared to mRNA expression. Levels of mRNA and protein correlate to a high degree during growth (across genes Spearman correlation = 0.65). The correlation decreases after the onset of development, mainly due to the time lag between mRNA transcription and protein translation. Hence, the data presented here show that the correlation between transcriptomics and proteomics is dependent on the conditions being studied and it is important to proceed with caution when using the transcriptome as a proxy for protein expression. The data presented in this study will also be a valuable resource for investigating *D. discoideum* development and are available in an interactive web app for ease of use: https://westholm.shinyapps.io/edelbroek_et_al_2024/.

## Results

### Experimental setup

The multicellular development of *Dictyostelium discoideum* starts when unicellular amoebae starve and embark on a developmental program. During this process, cells aggregate and go through distinct multicellular stages and culminate after 24 hours (h) in a fruiting body or “sorocarp” (Fig. 1). Here, we aimed to investigate how the transcriptome and proteome are regulated and correlated during early multicellular development of *D. discoideum*. Cells were starved on agar plates to induce multicellular aggregation, whereafter cells were harvested at time increments during 0 h to 10 h post starvation (Fig. 1). In order to minimize biological and technical variations, we collected cells from both halves of each plate and processed the cells for proteomics, and transcriptomics, respectively (Fig. 1).

**Fig. 1.**
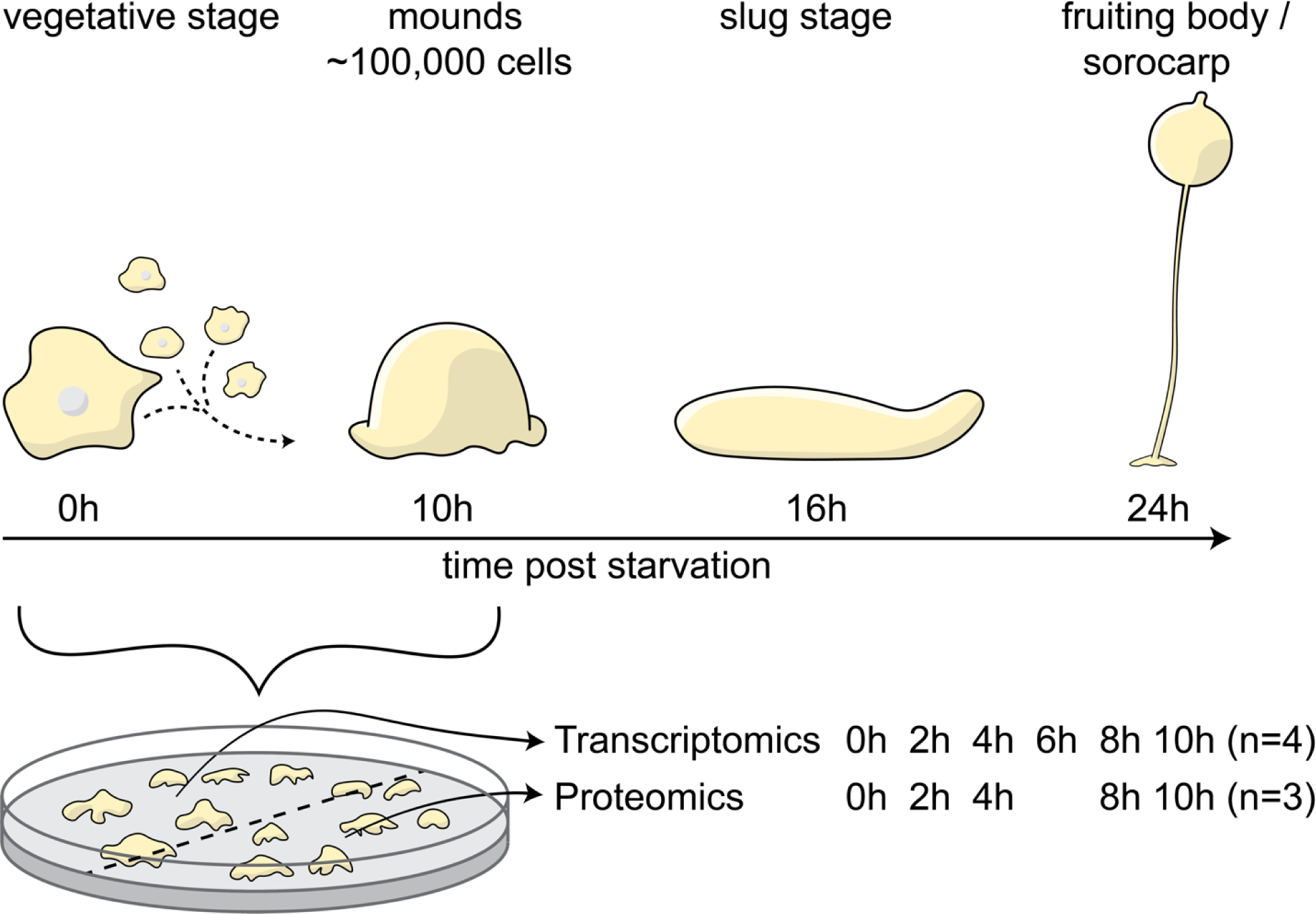
Experimental setup. Axenically grown cells were washed and plated on non-nutrient plates to induce multicellular development. Samples were taken from the same plate for transcriptomics and proteomics at 2h intervals, up to 10h post initiation of starvation.

Transcriptomics was performed on four biological replicates per time point for a total of 24 sequencing libraries. Proteomics was performed on three biological replicates and not from the 6h time point, resulting in 15 datasets.

### Major reorganization of the transcriptome during multicellular aggregation

The broad transcriptional changes over time were investigated by principal component analysis (Fig. 2a). Developmental time correlates highly with the first principal component, whereas the second principal component oscillates from 0h to 6h back to 10h. This is similar to what has been observed for cells developed on filters^15^. The biological replicates showed minimal variation, both in the PCA plot and by calculating their correlation (Fig. S1). From our data, we could identify 8310 protein coding transcripts differentially expressed (FDR-adjusted p-value <0.01) during the first 10h of development, suggesting that the great majority of the in total 11866 proteins are regulated at the transcript level during multicellular development (Table S1, 2).

**Fig. 2.**
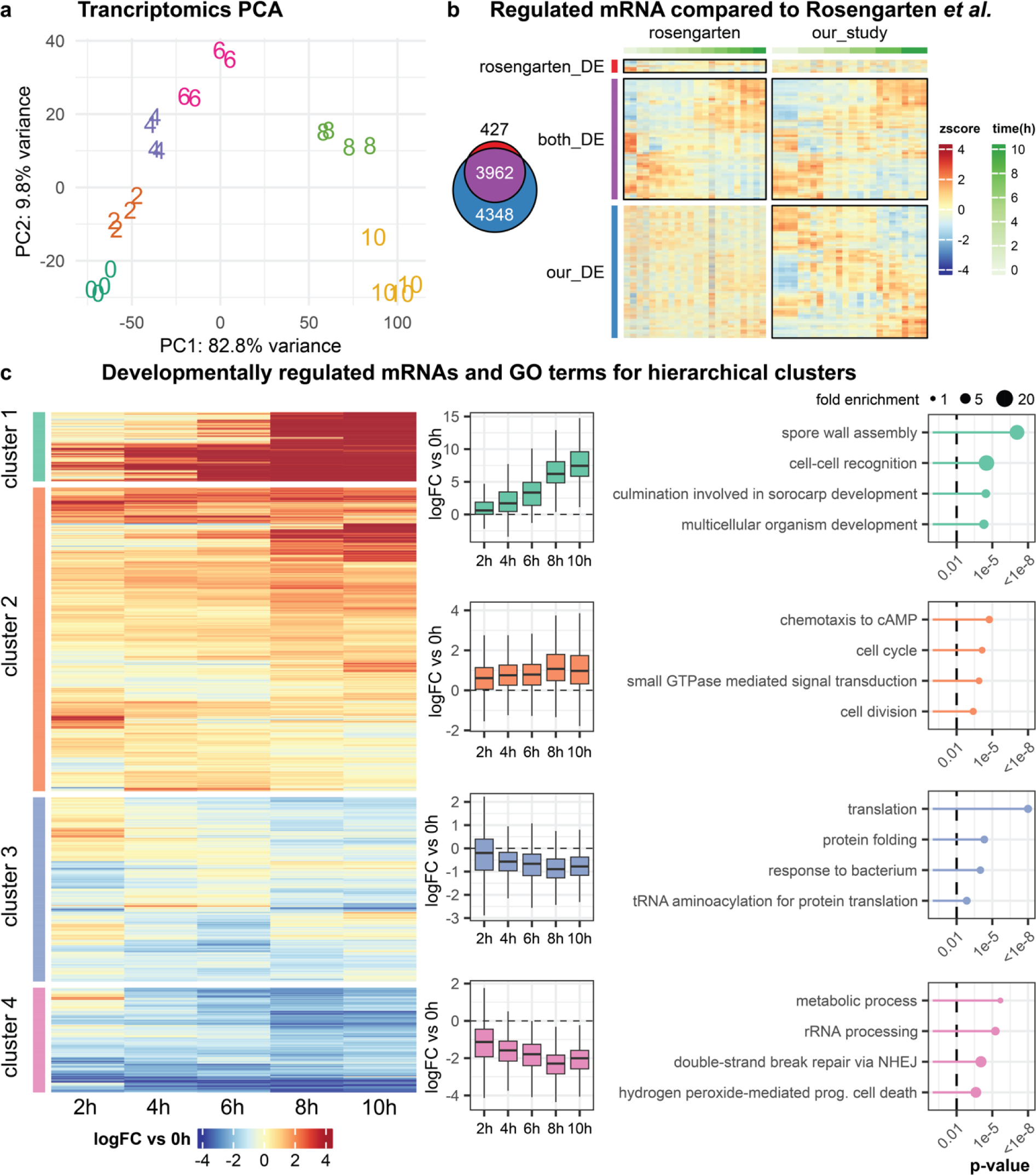
Major reorganization of transcriptome during multicellular aggregation. **a** Principal component analysis (PCA) of the developmental transcriptome based on the 500 transcripts that show the most variation. The first two principal components (PC1, PC2) are shown, which together explain 92.6% of the variance. Four biological replicates were analyzed per time point. Each replicate is plotted as a number, representing the time point of the replicate. **b** Comparison of the transcriptomics dataset generated in this study (our_study) with the 0h to 10h time points of the dataset generated by Rosengarten et al.^15^ (rosengarten). Red and purple: 3339 protein coding transcripts identified as differentially expressed in the Rosengarten dataset; blue and purple: 8310 protein coding transcripts in the dataset generated in this study; purple: 3001 protein coding transcripts identified in both studies. Regulation of the transcripts is shown by z-score from 0h growing cells to 10h post initiation of development, with differentially expressed transcripts outlined (black rectangles). **c** Hierarchical clustering of protein coding transcripts based on log fold change (logFC) versus the 0h time point. Transcripts were grouped into four main clusters, with the general regulation of each transcript shown in the heatmap to the left, and the general regulation of the cluster shown with boxplots for each time point to the right. The dashed line indicates a logFC of 0 versus the 0h time point. On the far right, the four most significant GO-terms for each cluster are shown, with Fisher’s exact test p-value for each GO-term (the dashed line indicates p-value 0.01). The size of the filled circles represents the fold enrichment of the GO-term in the cluster. For the full set of significant GO-terms, see Table S3

To compare our data with previous results from filter developed *D. discoideum* cells^15^, we re-analyzed their dataset and identified 4389 protein coding transcripts differentially expressed during the first ten hours of development (panels outlined with black square in Fig. 2b). Of these, 3962 overlapped with the differentially expressed genes identified in our experimental setup (Fig. 2b). The genes identified in both studies are generally regulated in the same manner during development (Fig. 2b).

The 8310 protein coding transcripts that were identified as differentially expressed were clustered based on their fold change relative to 0h at the different time points. Subsequently, the genes were split into four groups based on hierarchical clustering, i.e. highly upregulated in cluster 1, genes moderately upregulated in cluster 2, genes moderately downregulated from four hours of development in cluster 3, and genes strongly downregulated in cluster 4 (Fig. 2c). In order to classify the differentially expressed genes in each cluster, we performed Gene Ontology-terms (GO-terms) enrichment analysis. This showed that the highly upregulated genes in general are associated with processes connected to development of multicellularity, such as cell-cell recognition, culmination involved in fruiting body development, and assembly of the spore wall (ultimately leading to formation of spores) (Fig. 2c). Among the moderately upregulated genes, the formation of the multicellular aggregates is also represented, with terms corresponding to cAMP dependent chemotaxis and response to differentiation-inducting factor 1 (Table S3), but also the cell cycle is represented, with an enrichment of genes involved in DNA replication and cell division (Fig. 2c, Table S3). Terms in the clusters with downregulated genes include translation and metabolism. This is expected, since starvation induces growth arrest of the cells (Fig. 2c). In conclusion, the transcriptomics dataset describes multicellular aggregation in detail, and the GO-terms associated with differently regulated clusters aptly represent biological processes regulated during early development.

### The developmentally regulated proteome

In order to understand how well the transcriptomic data correlate with protein expression, we performed proteomic analysis, using cells from the same plates from which RNA was isolated for RNA-seq. By performing mass spectrometry (LC-MS/MS; label free quantification) on cells collected from several time points during early development (Fig. 1), 2478 proteins could be directly detected and quantified across all biological replicates and time points (Fig. 3a). For 1185 proteins, quantification was possible in all biological replicates at one or more time points, but was missing in replicates at other time points. We hypothesized that the lack of quantification in these replicates was mostly due to a lack of- or low expression of the protein at these timepoints. This is supported by the fact that proteins which lack quantification in some replicates also have a lower maximum expression level in the replicates with quantification (Fig. S2a). We therefore imputed missing values for these 1185 proteins, which were consistently expressed at a given time point, to avoid discarding them from analysis. The missing values were imputed using a probabilistic minimum, which accommodates for values missing due to low expression^18^. Addition of the imputed proteins brought the total number of proteins that we could analyze to 3663, about a third of all protein coding genes. For 7061 proteins no quantified peptides could be detected in the dataset. Many of these proteins are likely of low abundance or not expressed in accordance with the mRNA levels of these genes (Fig. 3a, Fig. S2b). Furthermore, the majority of unidentified proteins have a low annotation score, and low annotation quality may explain why some of the proteins were not identified (Fig. S2c).

**Fig. 3.**
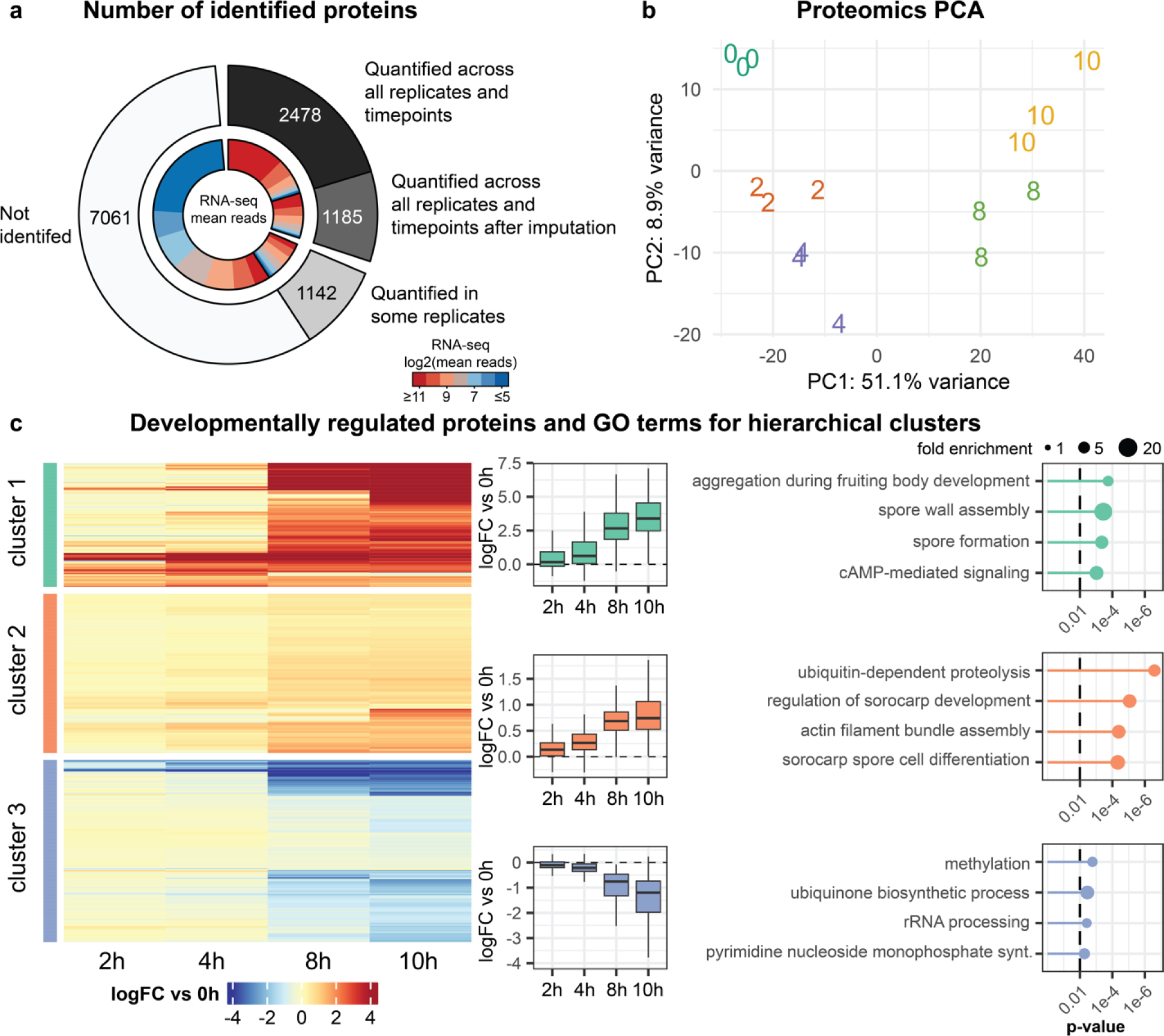
The proteome and its regulation during multicellular development. **a** Donut plot representing the total number of proteins, i.e. protein coding genes (outer circle) and the full transcriptome (inner circle). The inner ring shows the expression level from the transcriptomics analysis from high to low, and is correlated to each group of proteins. **b** Principal component analysis (PCA) of the proteomics dataset based on the 300 proteins that show most variation. The first two principal components (PC1, PC2) explain 60% of the variance in the dataset. The three replicates for each time point are plotted as separate numbers (time points in hours). **c** Hierarchical clustering of proteins based on log fold change (logFC) versus the 0h time point. The proteins were grouped into three main clusters, with the general regulation of each protein shown in the heatmap on the left, and the general regulation of the cluster shown with boxplots for each time point next to the heat map. The dashed line indicates a logFC of 0 versus the 0h time point. On the far right, the four most significant GO-terms for each cluster are shown, with the Fisher’s exact test p-value for each GO-term (the dashed line indicates p-value 0.01) and the size of the filled circle represents the enrichment of the GO-term in the cluster. For the full set of significant GO-terms, see Table S5.

Principal component analysis of the proteomics dataset resembles that of the transcriptomics and shows the same main trends, with the first principal component correlating with developmental time (Fig. 3b). Here too, variation between the biological replicates was low (Fig. S1). By analyzing the regulation of the quantified 3663 proteins, 672 were identified as differentially expressed during development (FDR-adjusted p-value <0.01) (Table S1, 4). In a study by Kelly e3/19/2024 5:42:00 AMt al., the *D. discoideum* proteome was analyzed during early multicellular development at 0.5h and 8h after initiating development in tissue-culture-treated plates^19^. In order to compare their findings with our proteomic analysis, we first reanalyzed their dataset. Subsequent comparison showed that the differentially expressed proteins from either dataset were regulated in similar manner over time (Fig. S3).

In the same way as for the transcriptomic analysis, we grouped the differentially expressed proteins from our study based on their fold change at different time points relative to the 0h time point (Fig. 3c). Interestingly, a distinct change can be observed at 8h of development where proteins are either upregulated (clusters 1 and 2) or down regulated (cluster 3). The first two clusters, which are made up of highly or moderately upregulated proteins, are associated with GO-terms linked to the development of multicellular aggregates and fruiting bodies, in line with what was observed in the transcriptomics analyses (Fig. 2c, Fig. 3c), but additionally protein ubiquitination and proteolysis appear to be upregulated (Table S5). The proteins that are downregulated from 8h are linked to growth arrest due to the lack of nutrients, where ribosomes and biosynthetic processes are broadly downregulated (Fig. 3c). Taken together, the proteomic dataset presented in this study describes a significant fraction of the total *D. discoideum* proteome during development, and is the first study to follow the regulation of the proteome during aggregative multicellularity in detail.

### High steady-state correlation of mRNA and protein levels

From the proteomics dataset it was possible to identify about one third of the total *D. discoideum* proteome across replicates and timepoints. 589 protein-coding genes are differentially expressed in both the transcriptomics and proteomics datasets. These were clustered according to their fold change relative to the 0h time point, and they appear to be largely regulated in the same manner during development (Fig. 4a, Fig. S4), illustrating that the mRNA and protein expression are be correlated.

**Fig. 4.**
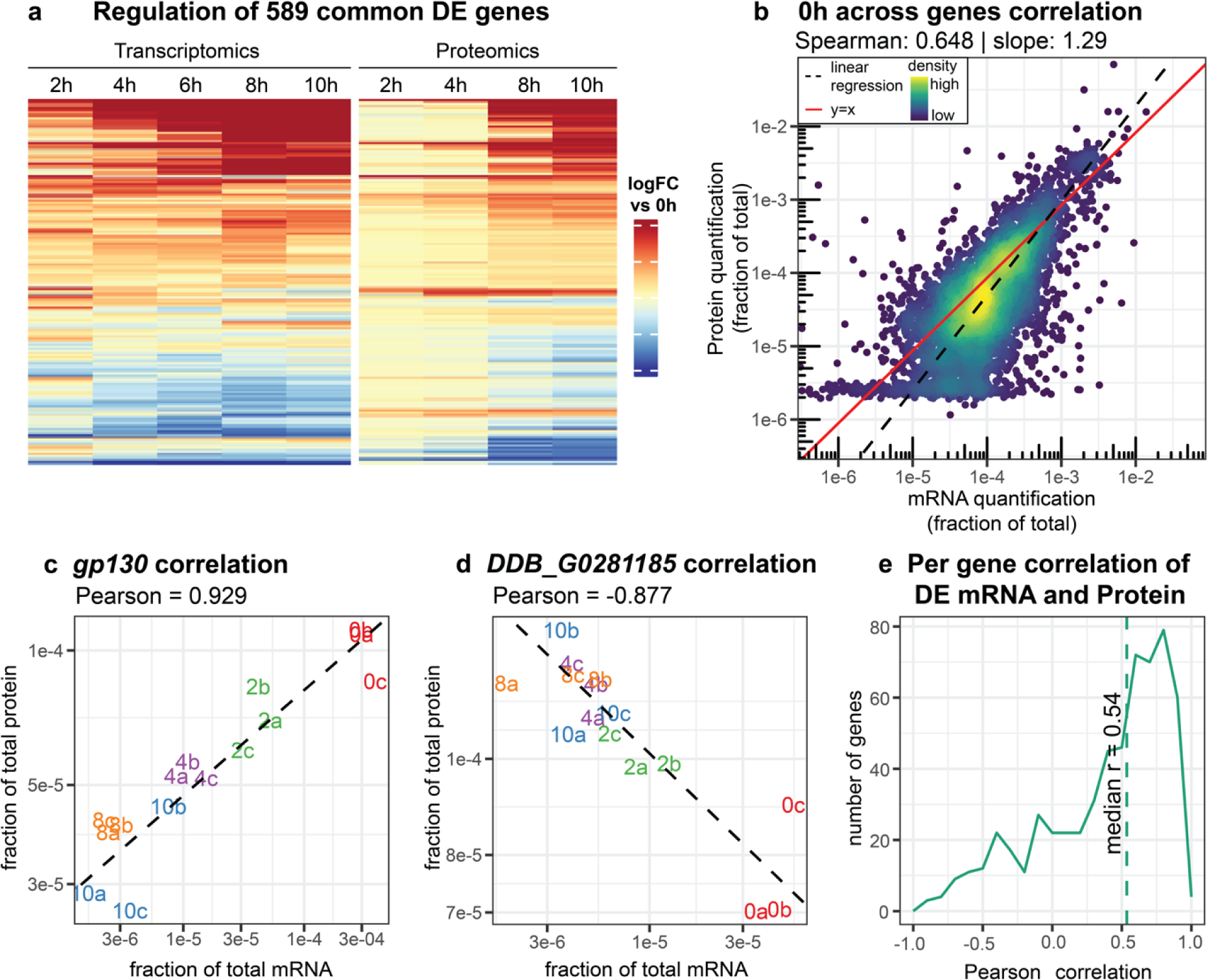
Correlation between mRNA and protein levels. **a** Regulation in log fold change (logFC) versus 0h time point for genes differentially expressed in both transcriptomics and proteomics datasets. The genes in the heatmap are hierarchically clustered based on their regulation. For an expanded plot with all genes differentially expressed in either dataset, see Fig. S4. **b** Correlation of the mean mRNA and protein levels across the 0h time point. Each dot represents the mean protein and mRNA expression from a single gene. The dashed black line indicates the linear regression of the data, the red line is the y=x diagonal. **c, d** Positive linear Pearson correlation of gp130, and negative correlation of DDB_G0281185, respectively. The expression in each sample is shown by fraction of total protein and fraction of total mRNA. Each biological replicate is plotted with a number, signifying the time point in hours, and letter, signifying the biological replicate. Only samples for which both transcriptomics and proteomics data was generated, are included. Linear regression is shown with a black dashed line, with the Pearson correlation above the plot. **e** Distribution of per gene Pearson correlations for all genes with differentially expressed mRNA and protein.

In order to study the correlation of the transcriptomic and proteomic datasets in more detail, we normalized the expression of each gene by the quantification of all genes, such that all mRNA values or protein values at a given time point sum up to 1. Across all genes at the 0h time point, the mRNA levels and protein levels correlated well (Spearman correlation 0.65, Fig. 4b. At later time points however, the correlation dropped to 0.56 (Fig. S5). It should also be noted that at all time points, the regression of the data had slope greater than 1 (Fig. 4b Fig. S5). This is indicative of the fact that distribution of the data is different between the transcriptomics and proteomics datasets. We see, relative to the mRNA abundance, a wider dynamic range as well as a more uneven distribution of protein expression levels (Fig. S6). For example, for genes quantified in both datasets, the top 2 proteins together encompass 10% of the total protein abundance, whereas the top 2 most abundant transcripts encompass a more modest 2.3%. A slope greater than 1 in the mRNA-protein correlation has previously been observed for human tissues as well as in yeast^20,21^.

Although the mRNA and protein levels correlated well for each time point, we wondered if this is also the case for the regulation of each of these molecules over all samples. For instance, if an mRNA is upregulated at a specific time point during development, does this also hold true for its cognate protein? As an example, the *gp130* mRNA and its associated glycoprotein 130 are both downregulated during development, resulting in a high correlation (Pearson’s r = 0.93, Fig. 4c). For other genes, a decrease in mRNA abundance between samples coincided with increased protein abundance resulting in a negative correlation (Fig. 4d). When considering the Pearson correlation of all genes that were differentially expressed in both the transcriptomics and proteomics datasets, we observed a positive median correlation (median Pearson’s r = 0.54, Fig. 4e, Table S6). Hence, in most cases an upregulation of mRNA corresponds to an increase in protein, and vice versa, for all differentially expressed genes. Notably, the median correlation is much lower when considering genes which are only differentially expressed in one of the datasets, or not differentially expressed at all (median Pearson’s r = 0.16, Fig. S7a). By calculating the correlation per gene, it is also possible to show the advantage of our sampling setup – to isolate mRNA and protein from the same plate (biological replicate) (Fig.1). When we compare to mismatched samples, i.e. when mRNA from one replicate (plate) is compared to protein from another replicate, for the same time point, the median correlation was significantly lower as compared to matching samples from the same plate (Fig. S7b, c).

In sum, mRNA and protein expression are in general well correlated. Across genes, correlation is highest during steady state growth. The median per gene correlation is high for differentially expressed genes, but not for genes which lack regulation in the transcriptome or proteome during development.

### Differences between mRNA and protein regulation

To better understand the details behind the expression patterns in the transcriptomics and proteomics data, we performed an integrated unsupervised analysis using MEFISTO^22^. MEFISTO is a factor analysis method that reduces a multi-omics data set into a few latent factors that explain most of the variance in the full data set. Running MEFISTO on all genes for which we have both protein and mRNA data (Table S1), resulted in three factors that explain most of the variance in the transcriptomics data, and out of these three factors, Factor 1 also explained a large fraction of the variance in the proteomics data (Fig. 5a). Factor 1, which makes up 25% of the variance in the transcriptomics data and 34% in the proteomics data, represents steadily increasing or decreasing expression over the developmental time course (Fig. 5b). Factor 2, explaining 27% and 3% of the variance in the transcriptomics and proteomics data, respectively, represents a pattern where expression decreases between 0 and 4 hours, after which it plateaus and then increases at 8 and 10 hours, or vice versa. Factor 3, which explains 17% and 6% of the variance respectively, shows a dramatic increase or decrease in expression between 0 and 2 hours, after which the expression gradually returns (Fig. 5b). This shows that mRNA is more dynamically regulated, with more variable expression patterns, whereas proteins mostly show steadily increasing or decreasing levels over the developmental time course.

**Fig. 5.**
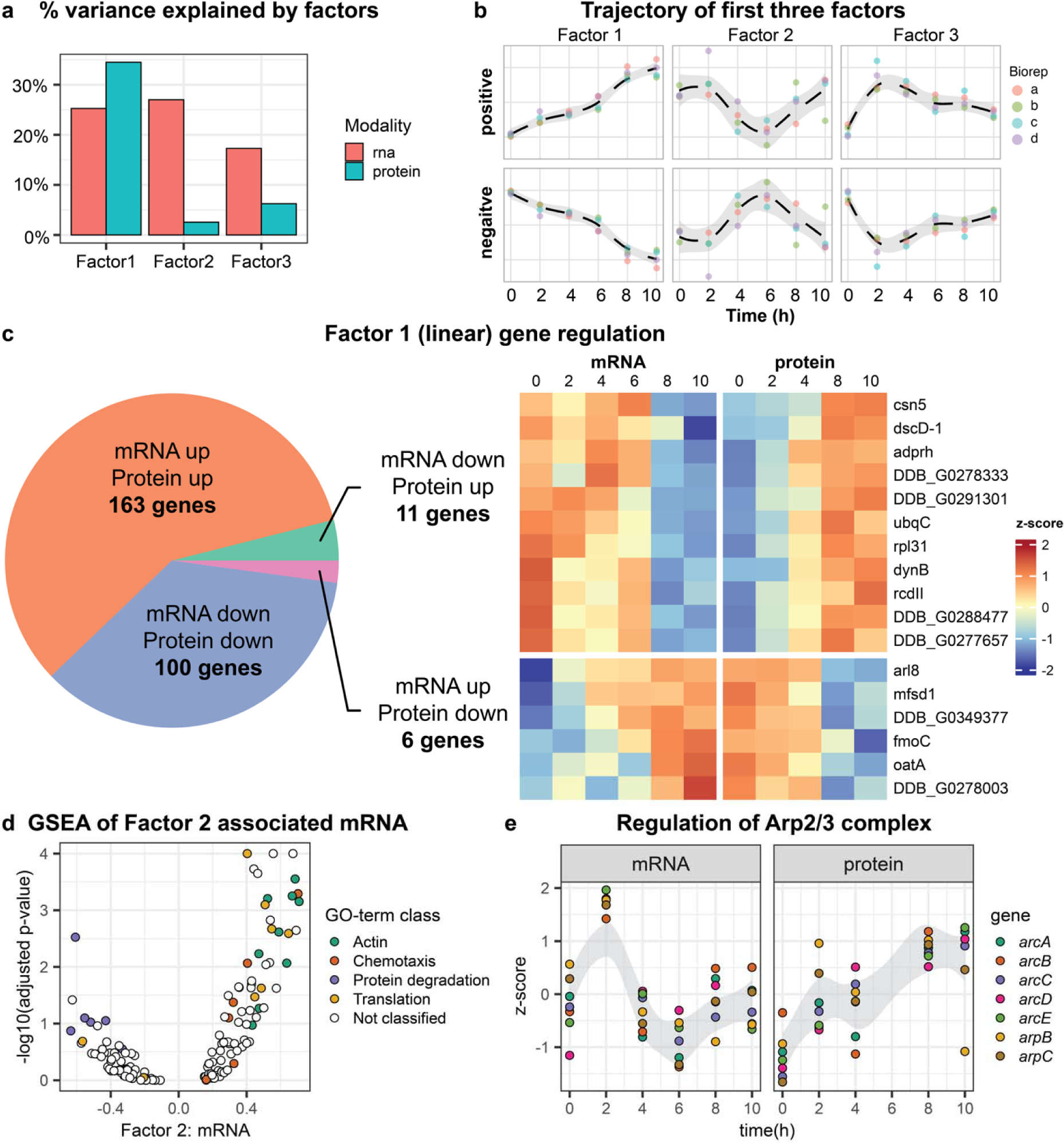
Multi-omics factor analysis of mRNAs and proteins. **a** Percentage of variance explained by the first three factors of the multi-omics factor analysis for the transcriptomics and proteomics datasets. **b** Trajectory of the first three factors over time. The different replicates are shown, and the dashed line is the loess (locally estimated scatterplot smoothing) regression through the replicates. **c** Analysis of genes with high Factor 1 values at both mRNA and protein modalities.

For 280 genes, both the mRNA and protein were highly associated with Factor 1, i.e. linearly up- or downregulated during development. The majority of these were regulated in the same direction in the mRNA and protein modalities (Fig. 5c). For 11 genes however, mRNA was linearly downregulated and protein upregulated, and vice-versa for 6 other genes (Fig. 5c).

Since the majority of the mRNA regulation could be explained not by a steady increase or decrease but by more dynamic patterns, we investigated what GO-terms are associated with the dynamically regulated mRNA Factor 2 values (Fig. 5d). Genes with high Factor 2 values are upregulated at the 2h time point, downregulated from 4h-8h, followed by upregulation at the 10h time point. Among these genes, GO-terms related to translation and actin are enriched (Fig. 5d). This is reflected in regulation of the Arp2/3 complex, a major regulator of the actin cytoskeleton. At the mRNA level, the regulation is similar to Factor 2 (Fig. 5e). On the other hand, the complex appears to be linearly upregulated at the protein level, matching Factor 1 (Fig. 5e). The proteasome complex on the other hand, is associated with negative Factor 2 values (mRNA), but is also linearly upregulated (protein) (Fig. S8a, b).

In conclusion, the factor analysis revealed several major trajectories of the dynamic mRNA regulation, which all appear to be paired with linear up- or downregulation at the protein level.

The proportion of genes with different regulations shown in the pie-chart. Genes with opposing mRNA and protein regulation are represented in the heatmap. **d** Gene set enrichment analysis (GSEA) of Factor 2 mRNA values, with GO-terms classified under broad terms. The x-axis shows the average Factor 2: mRNA values while the y-axis shows the GSEA p-values, after negative log transformation. **e** Regulation of members of the Arp2/3 complex. For each gene, z-scores were calculated from the mean expression per time point. The gray zone denotes the 95% confidence interval trajectory of all z-scores.

### Protein expression is generally delayed several hours compared to mRNA expression

From the previous analyses, we identified that for a number of genes, the mRNA and protein regulation opposed one another (Fig. 5c, d). We hypothesized that for some of these genes, the difference may be explained by a time lag between the mRNA transcription and protein translation. To investigate this, we calculated the Spearman correlations of mRNA and protein levels, as before (Fig. 4a, Fig. S5), but matched all of the transcriptomics time points with all of the proteomics time points. Matching mRNA and protein expression from the same time points shows Spearman correlations of 0.65 to 0.56, however, for all time points, we found higher correlations when matching the mRNA expression with protein expression 2 to 4 hours later (Fig. 6a). This observed time lag is in agreement with previous studies in yeast^23^ and *Drosophila*^24^. When exclusively considering genes that are differentially expressed during development according to the protein and mRNA expression, the trend is largely the same (Fig. 6b). Here, however, the maximum correlation is higher, similar to what we previously observed (Fig. 4b). Since these are genes which are highly affected throughout development, there is a larger difference between highly correlated pairs of time points (e.g. mRNA at 0h vs proteins at 4h, Spearman correlation = 0.71, Fig. 6b) and those that show low correlation (e.g. mRNA at 10h vs proteins at 0h protein, Spearman correlation = 0.02, Fig. 6b).

**Fig. 6.**
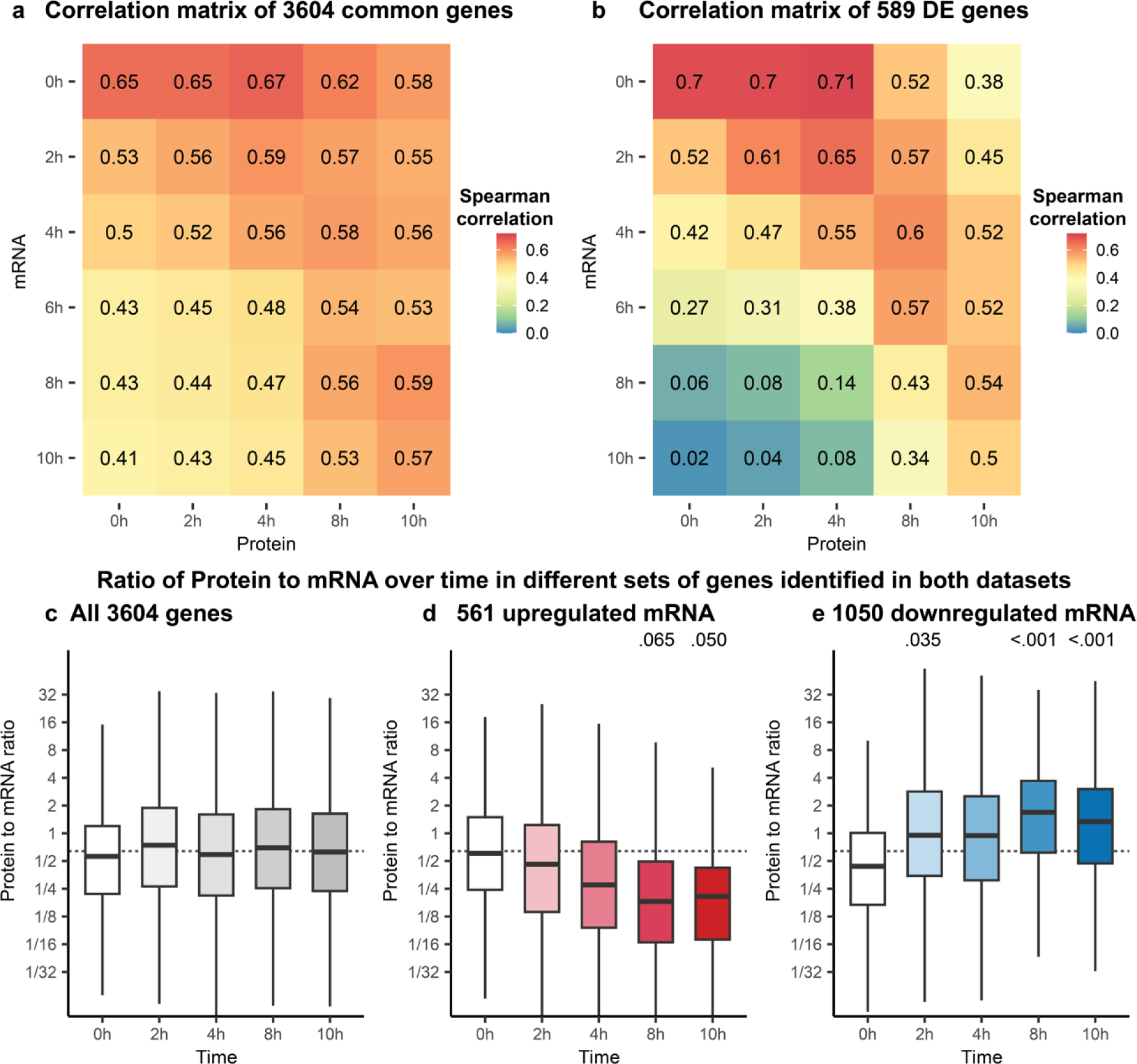
Time lag between mRNA and protein. **a,b** Spearman correlations of mean protein values and mean mRNA values across genes for time points from 0h to 10h. In **a**, all genes quantified in both datasets are included in the analysis. In **b**, only differentially expressed (DE) genes are included. **c-e,** Ratio of protein to mRNA for different time points. Boxplots are based on **c**: all identified in both datasets; **d**: genes for which the mRNA was upregulated at 10h; **e**: genes for which the mRNA was downregulated at 10h. Above the boxplots: Dunnett contrasts p-values relative to 0h time point reported for time points with p-values lower than 0.1. Dashed lines indicate the median protein to mRNA ratio for all genes.

Next, we investigated if the ratios of protein to mRNA are affected during multicellular development. By dividing the protein quantification by the mRNA quantification, the protein to mRNA ratio could be calculated for each gene, at each time point. It should be noted that these ratios are in no way indicative of the absolute numbers of protein or mRNA molecules, and are purely relative values. When considering all genes expressed in both omics datasets, there are no significant differences in protein to mRNA ratios at the different time points (Fig. 6c). There are however significant differences between timepoints when considering genes for which the mRNA is either up- or downregulated during multicellular development (Fig. 6d, e, Fig. S9). 10h after onset of multicellular development, the ratio of protein to mRNA is significantly decreased for genes which are upregulated (mRNA) (Fig. 6d). In contrast, downregulated genes show the opposite effect, with ratios of protein to mRNA significantly increasing over time (Fig. 6e). We suspect that this is largely another effect of the time lag between mRNA transcription and protein translation. Those genes that are upregulated at the transcriptional level have a relatively lower level of protein until translation catches up or the mRNA is eventually downregulated again. The opposite is true for genes which are downregulated, here the protein needs to be turned over for the levels to agree with the mRNA.

Taken together, these results show a modest correlation between protein and mRNA levels analyzed at the same developmental time points, however, the correlation increases when considering protein samples taken 2-4 hours after the mRNA samples.

## Discussion

The evolution of multicellularity is thought to have occurred several times, through both clonal mechanisms, such as in animals and plants, and through aggregative mechanisms, where cells stream together to form multicellular structures^25,26^. Aggregative multicellularity has been studied using the social amoebae, where processes behind the transition from uni- to multicellular life have been investigated^27–29^. Numerous studies that focused on individual genes, have identified some of the key players involved in this transition^30^, but in recent years next-generation sequencing methodology have paved the way to acquire a complete understanding of this process at the transcriptional level^13–17^.

Here, we report transcriptomic and proteomic analyses of *D. discoideum* during early multicellular development. Using LC/MS-MS, we were able to capture the regulation of roughly a third of the proteome during development. This was combined with transcriptomic analyses, covering the great majority of the genes. By analyzing the two data sets separately, but also combining them with factor analysis, we identified several biological processes, vital for *D. discoideum* multicellular development. Together, this provides us with a detailed picture of gene expression and regulation, from mRNA to protein, during early development of the social amoeba.

Included in the upregulated processes, we find chemotaxis and development of the sorocarp, whereas ribosome biogenesis and general metabolism are among the most downregulated (Fig. 2c, Fig. 3c), in line with what has been observed previously^15^. Interestingly, some processes were dynamically up- and downregulated at the mRNA level, but linearly regulated at the protein level. This is similar to what has been observed in vertebrates, where a spike or dip in mRNA causes a switch in protein levels that are either down- or upregulated^10,31^. Examples of this kind of regulation in *D. discoideum* are protein degradation processes and actin related genes, illustrating that relying on mRNA levels only for insights into the temporal impact of these processes may be misleading. Notably, for the majority of regulated mRNAs in our dataset, the protein response is delayed, and the proteins emerge first about 2h to 4h after mRNA appearance (Fig. 6). This result can likely explain the previous observation that early major morphological changes do not coincide with transcriptomic data, i.e. the phenotypes connected to the expressed mRNAs are delayed^13,15^. This result demonstrates that the proteome more accurately describes the functional gene expression and resulting phenotype^6^. Hence, it may be preferable to rely on proteomics, and not transcriptomics, for assigning genes to a specific temporal phenotype or morphological stage.

To what extent mRNA and protein expression correlate in different organisms remains largely unknown. For a solid comparison, the data should be generated from the same original sample, and should contain minimal technical variability^5^. In our study, we could verify that technical variation was very small and observed a significant increase in correlation due to the sampling approach (Fig. S1, Fig. S7c). At the 0h time point, prior to development of the cells, we observed a Spearman correlation across genes of 0.65 (Fig. 4a). This is somewhat lower than what has been recently reported for bacteria^32^ (Spearman = 0.80) and more in line with reports of mammalian cells and other eukaryotes^6^. Maybe this is reflective of a more linear relationship between mRNA and protein in bacteria than in eukaryotes. Additionally, some of the dissimilarity might be due to different methods or genes selected for comparison. For example, we observed Spearman correlations as high as 0.70 when considering only differentially expressed genes.

Besides the genome wide correlation between mRNA and protein levels at distinct time points, we also investigated the correlation between mRNA and protein for individual genes across all time points (Fig. 4b). Here, however, we found a relatively low median Pearson correlation of 0.16. One reason for this low correlation is that while the majority of the transcriptome is regulated during development (Fig. 2b, Fig. 5a), only a fraction of the proteome was clearly developmentally regulated. Thus, if we restrict our analysis to genes that were differentially expressed in both transcriptomic and proteomic datasets, we observe a drastically increased median correlation of 0.54 (Fig. 4b). This is similar to what was observed in a xenograft model^33^. Another factor contributing to the low correlation between individual mRNAs and their corresponding proteins over time, is the time-lag discussed above.

To conclude, the data presented here enable in-depth study of aggregative multicellularity at both transcript and protein levels, and can constitute a significant resource for comparative studies of other members of Amoebozoa. Notably, we show that the overall correlation between mRNA and protein in *D. discoideum* at steady-state is rather high, but correlations of individual genes vary, and care should be taken when inferring the presence of proteins from transcriptomic data. We are pleased to refer anyone interested to explore the RNA-protein expression during early development in *D. discoideum* to the easy-to-use interactive web application https://westholm.shinyapps.io/edelbroek_et_al_2024/.

## Methods

### Growth conditions

*D. discoideum* AX4 wildtype cells (DictyStockCenter ID: DBS0237637) were grown axenically in HL5-C (Formedium) to exponential phase. 3×10^8^ cells were harvested at 400 x g, 5 min, and washed twice in 50 ml KK2 (2.2 g/l KH_2_PO_4_, 0.7 g/l K_2_HPO_4_). For the 0h time point (not developed), half the cells were harvested as before and stored at -80°C for subsequent processing for transcriptomics library preparation; the other half was harvested and stored at - 80°C and later used for proteomics sample preparation. For the other time points, cells were plated on 92mm NN-Agar plates (1.2 g/l KH2PO4, 0.48 g/l Na2HPO4⋅2H2O, 15 g/l agar) and harvested at the defined time points using Nunc Cell Scrapers (Thermo Fisher), into KK2 buffer; half the plate for transcriptomics and half the plate for proteomics. The cells were treated as described above and the cell pellets were frozen at -80°C until further processed for transcriptomics or proteomics sample preparation.

### Transcriptomics library preparation and sequencing

The frozen cell pellets were dissolved in 1 ml TRIzol Reagent (Invitrogen) and total RNA was prepared according to the user guide, except with an additional 75% EtOH wash of the RNA pellet. Following RNA extraction, 15ug total RNA samples were DNase treated using TURBO DNase (Invitrogen) according to manufacturer’s protocol and purified by phenol/chloroform extraction. 75 μl Phenol stabilized: Chloroform: Isoamyl Alcohol (25:24:1, PanReacAppliChem) was added to 75 μl DNase treated RNA, shaken for 20 s and centrifuged 5 min, 16 000 x g. The upper phase was transferred to new tubes with 187.5 μl EtOH (99%), 7.5 μl 3M Sodium Acetate, 5 μg glycogen, and the RNA was precipitated at -20°C overnight. The RNA was harvested (16 000 x g, 30 min, 4°C), washed with 150 μl 75% EtOH (16 000 x g, 10 min, 4°C), and resuspended in 50 μl RNase free H_2_O. Sequencing libraries were prepared from 700 ng total RNA using the TruSeq stranded mRNA library preparation kit (Cat# 20020594/5, Illumina Inc.) including polyA selection. The library preparation was performed according to the manufacturers’ protocol (#1000000040498). Libraries were sequenced on the NovaSeq 6000 System (Illumina) on two SP Flowcells, with single reads, 100bp read length (v1 chemistry).

To enable mapping of the sequencing reads, adapters were trimmed using cutadapt v2.10^34^. Trimmed reads from different sequencing lanes were pooled and mapped using STAR v2.7.5, allowing a maximum intron size of 2000 bases^35^. Mapped reads from both Flowcells were merged with samtools v1.10^36^. Reads were assigned to genes with featureCounts, part of the subread v2.0.1 package^37^. For both read mapping and counting, the improved *D. discoideum* gene annotation was used^38^. mRNA library preparation and sequencing were performed at SciLifeLab Uppsala.

### Proteomics sample preparation and LC-MS/MS analysis

The cell pellets were lysed in 150 µL of 1% β-octyl glucopyranoside and 6M urea containing lysis buffer using a sonication probe for 60 seconds (3 mm probe, pulse 1 s, amplitude 30%) according to a standard operating procedure. After homogenization, the samples were incubated for 90 min at 4°C during mild agitation. The lysates were clarified by centrifugation for 10 min (16 000 × g at 4°C). The supernatant containing extracted proteins was collected and further processed. The total protein concentration in the samples was measured using the DC Protein Assay (BioRad) with bovine serum albumin as standard. Aliquots corresponding to 35 µg of proteins were withdrawn for digestion. The proteins were reduced, alkylated, and on-filter digested by trypsin using 3kDa centrifugal spin filter (Millipore, Ireland). The collected peptide filtrate was vacuum centrifuged to dryness using a Speedvac system. The samples were dissolved in 100 µL 0.1% formic acid and further diluted 4 times. For LC-MS/MS analysis, the peptides were separated in reversed-phase on a C18-column with 150 min gradient and electrosprayed on-line to a Q Exactive Plus Orbitrap LC-MS/MS system (Thermo Scientific). Tandem mass spectrometry was performed applying Higher-energy collisional dissociation.

Label free quantification (LFQ) of the raw data was performed using FragPipe v20.0 (https://fragpipe.nesvilab.org/), which is powered by MSFragger^39^. Analysis was performed with oxidation and lysine ubiquitination specified as variable modifications. Up to 3 missed cleavages were allowed. PSM validation performed with Percolator^40^, and protein inference with ProteinProphet^41^. Data is filtered at 1% FDR at the PSM, ion, peptide, and protein levels. Site localization with PTM-Prophet. For quantification, a minimum of 1 ion was required for MaxLFQ determination with IonQuant, using match between runs^42^.

### RNA-seq analysis

Counts from transcripts encoding the same protein were summed, and transcripts not encoding proteins were discarded, to allow for analysis of protein-coding transcripts and enable downstream comparison to protein data. Differentially expressed genes over time from mRNA-seq were identified with DESeq2 v.1.41.12, using a likelihood ratio test to compare a model where gene expression is explained by developmental time to a null model of constant expression^43^. Genes with an FDR-adjusted p-value below 0.01 were designated as differentially expressed. Normalized, transformed count data was extracted using variance stabilizing transformations, and shrunken log fold changes of differentially expressed genes were calculated with Approximate Posterior Estimation for generalized linear model^44^. Counts from 0h to 10h time points from Rosengarten *et al.*, were processed in the same manner ^15,45^. Genes plotted in heatmaps were hierarchically clustered based on their log fold changes or z-scores. Gene set enrichment for GO-terms was performed using topGO with the *weight01* algorithm and using Fisher’s exact test to determine statistical significance.

### Proteomics analysis

Protein quantification is based on MaxLFQ values from FragPipe. Values were imputed for proteins that were quantified in all biological replicates at a given time point, but where values were missing at other time points. Imputation was performed using a probabilistic minimum from the imputeLCMD v.2.1 R package^18^. Differentially expressed proteins (FDR-adjusted p-value 0.01) were identified with Limma v.3.57.11 by fitting linear models, with empirical Bayes smoothing^46,47^. The data by Kelly et al.^19,48^, was imputed and processed identically for comparison of differentially expressed genes. Clustering and GO-term analysis was performed as for the mRNA data.

### Integrative analysis

Genes which were quantified in both the transcriptomics and proteomics datasets, were utilized for integrative analysis. To enable comparison of the mRNA and protein levels, the values of each replicate, for each dataset, were divided by the total sum of values for that replicate such that the scaled values sum to 1. For across genes correlation at a single time point, the mean mRNA and protein levels were calculated from the biological replicates. Linear regression was calculated with ranged major axes using lmodel2 v.1.7.13. For per-gene correlations, Pearson correlations were calculated for each gene with all replicates available from both transcriptomics and proteomics.

Multi omics factor analysis was performed with MOFA v.1.11.0 using data from all time points^22^. For analysis based on Factor 1, genes were selected with a Factor 1 loading at both mRNA and protein modalities above 0.45 or below -0.45. Gene set enrichment analysis of Factor 2 mRNA was performed based on the Factor 2 gene weights using piano v.2.17.0^49^.

For time lag analysis, the Spearman across-genes correlation was calculated for each transcriptomics time point with each proteomics time point, either with all common quantified genes, or those that were differentially expressed in both datasets. To calculate the ratios of protein levels to mRNA levels, the normalized protein level was divided by the normalized mRNA level for each gene. Differentially expressed genes at the mRNA modality, which have a fold change above 2 at the 10h time point compared to the 0h time point, were identified as upregulated, and those with a fold change below 0.5 as downregulated.

## Data availability

Complete proteomics data submitted to MassIVE, with accession number MSV000093620, and is linked to ProteomeXchange: https://doi.org/10.25345/C5H12VJ75. The transcriptomics dataset of all 24 sequencing libraries has been submitted to GEO with accession number GSE249880.

## Code availability

All code for downstream analysis of the transcriptomics and proteomics datasets can be accessed at https://doi.org/10.6084/m9.figshare.25365283 together with the generated figures and tables.

## Acknowledgements

Sequencing was performed by the SNP&SEQ Technology Platform in Uppsala. The facility is part of the National Genomics Infrastructure (NGI) Sweden and Science for Life Laboratory. The SNP&SEQ Platform is also supported by the Swedish Research Council and the Knut and Alice Wallenberg Foundation. Proteomics was performed at the MS-based proteomics facility platform at Uppsala University. Dr. Anna Widgren is acknowledged for her support with the analysis. We would like to thank Jonas Kjellin and Johan Reimegård for critical reading of the manuscript. The computations and data handling were enabled by resources provided by the National Academic Infrastructure for Supercomputing in Sweden (NAISS) at Uppsala University, partially funded by the Swedish Research Council through grant agreement no. 2022-06725. Jakub Orzechowski Westholm is financially supported by the Knut and Alice Wallenberg Foundation as part of the National Bioinformatics Infrastructure Sweden at SciLifeLab. This work was supported by grant no. 2021-05793 from Swedish Research Council (Vetenskapsrådet) and grant no. CTS 18:381 from Carl Tryggers Stiftelse to Fredrik Söderbom.

## Author contributions

BE, JOW and FS designed the study. BE grew cell cultures and prepared samples for transcriptomics and proteomics. JB led the proteomics data generation. BE and JOW analyzed the data and prepared figures. BE, JOW and FS drafted the manuscript. All authors read and approved the final manuscript.

