## Supplemental Information for "Multi-omics analysis of aggregative multicellularity"

### Supplemental Figures

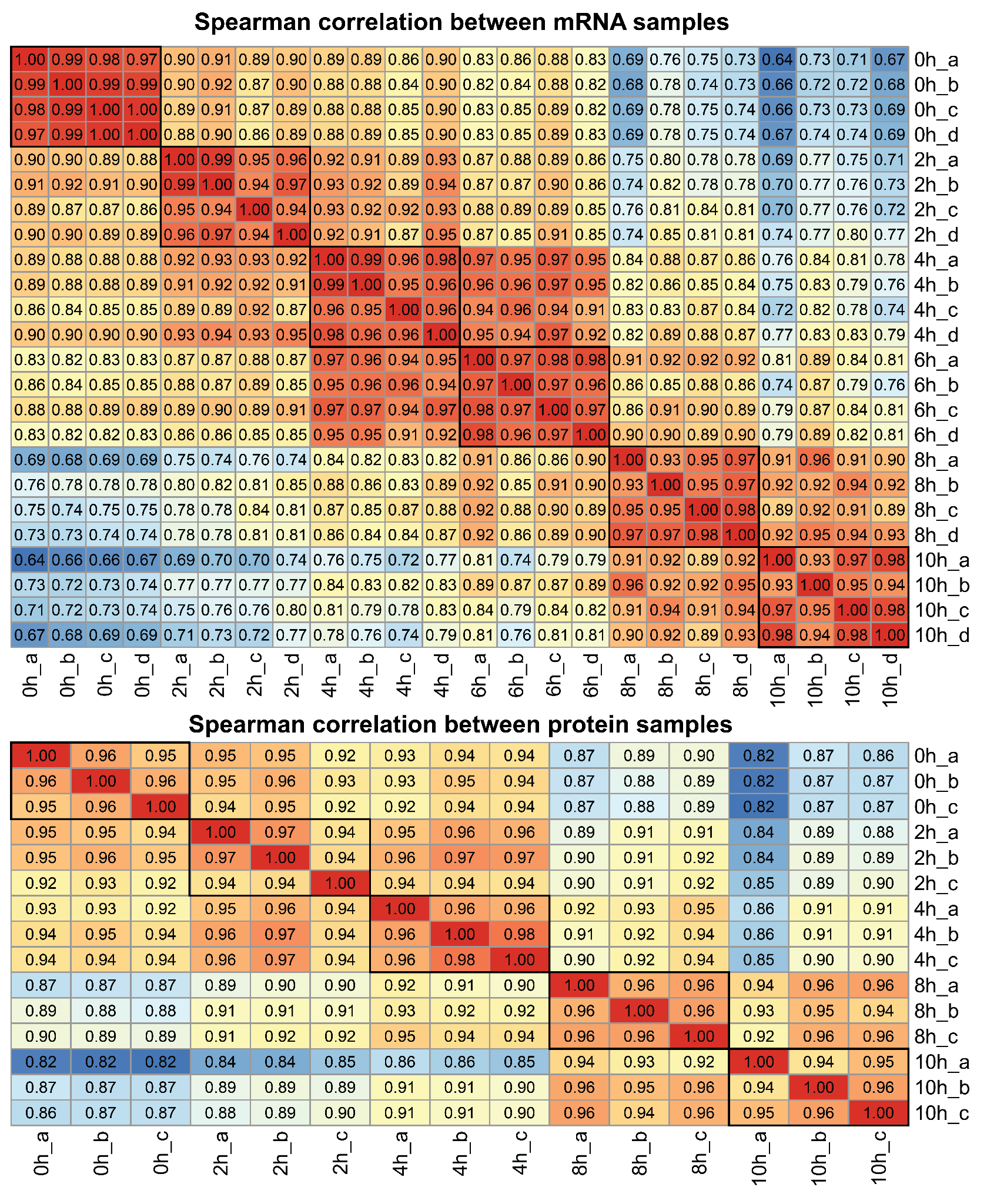

**Fig. S1 | Correlation between biological replicates of transcriptomics and proteomics.** Correlation matrix of all transcriptomics samples (top) and proteomics samples (bottom), with the Spearman correlation indicated for each comparison. Biological replicates of the same time point are indicated with a black box.

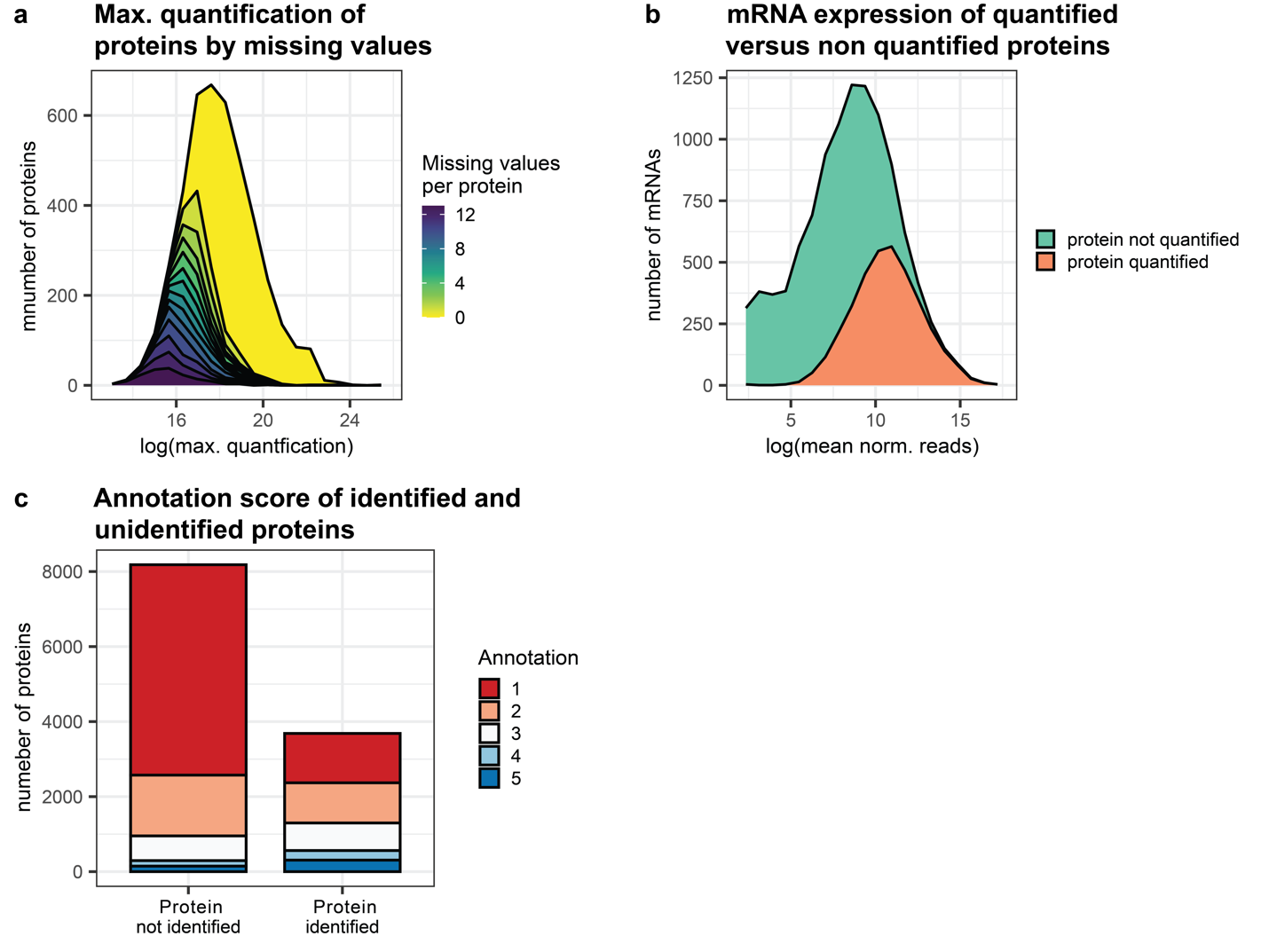

**Fig. S2 | Analysis of missing proteins. a** Protein quantification grouped by the number of missing values. For each protein, the maximum value among all samples was calculated. Proteins without missing values across all samples are yellow. For proteins with all missing values, no maximum value could be calculated and these are omitted. **b** mRNA expression in log-scaled normalized reads. Reads were averaged across biological replicates and time points for each gene, classified by whether or not the cognate protein was quantified from the proteomics analysis. **c** Annotation score of not identified proteins, and identified proteins, ranging from 1 (lowest annotation score) to 5 (maximum annotation score), accessed from UniProt^1^.

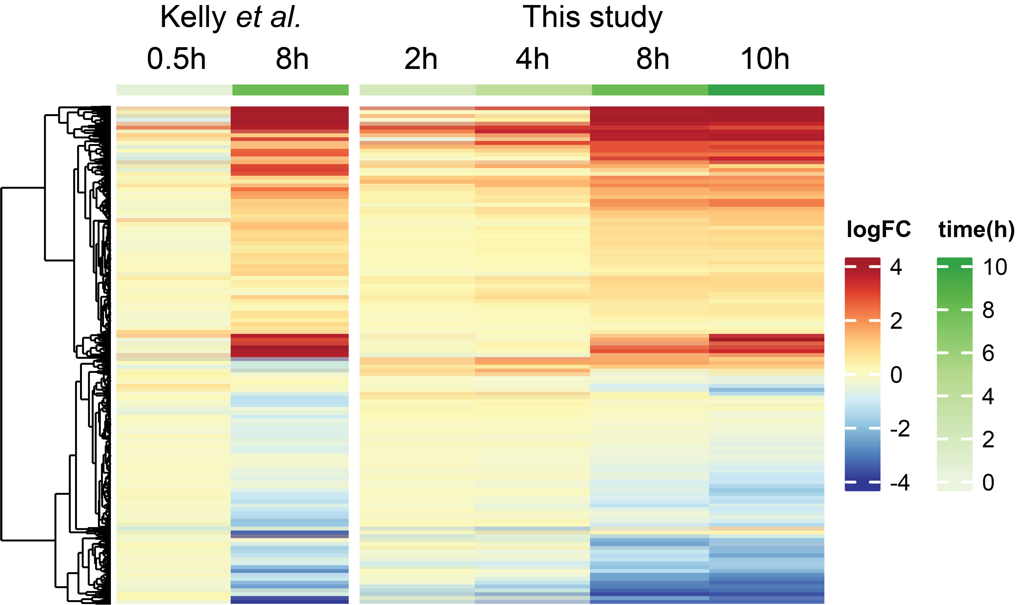

**Fig. S3 | Regulated proteins compared to dataset by Kelly et al.** Dataset was accessed through the ProteomeXchange ^2,3^. Regulation in log fold change (logFC) of time points relative to non-developed cells. Proteins included in heatmap were differentially expressed in either dataset. Proteins with missing values in either dataset, were omitted.

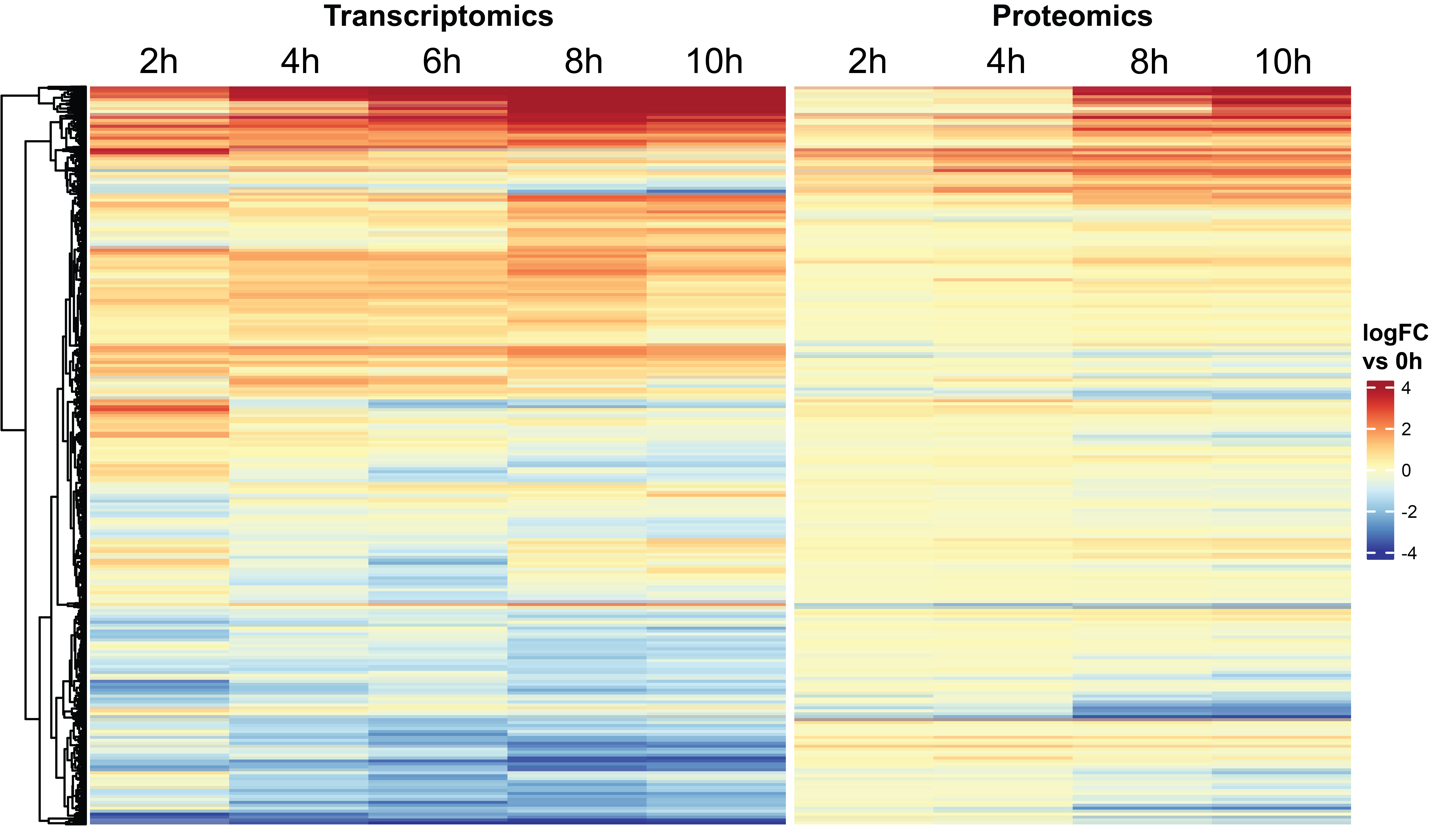
**Fig. S4 | Regulation of genes for both transcriptomics and proteomics datasets for all differentially expressed genes.** All genes that are differentially expressed in either dataset, and contained no missing values, are plotted (2934 genes). Log fold change (logFC) is calculated for the indicated time points versus the 0h time point. The genes in the heatmap are hierarchically clustered based on their regulation, with the dendrogram shown left.

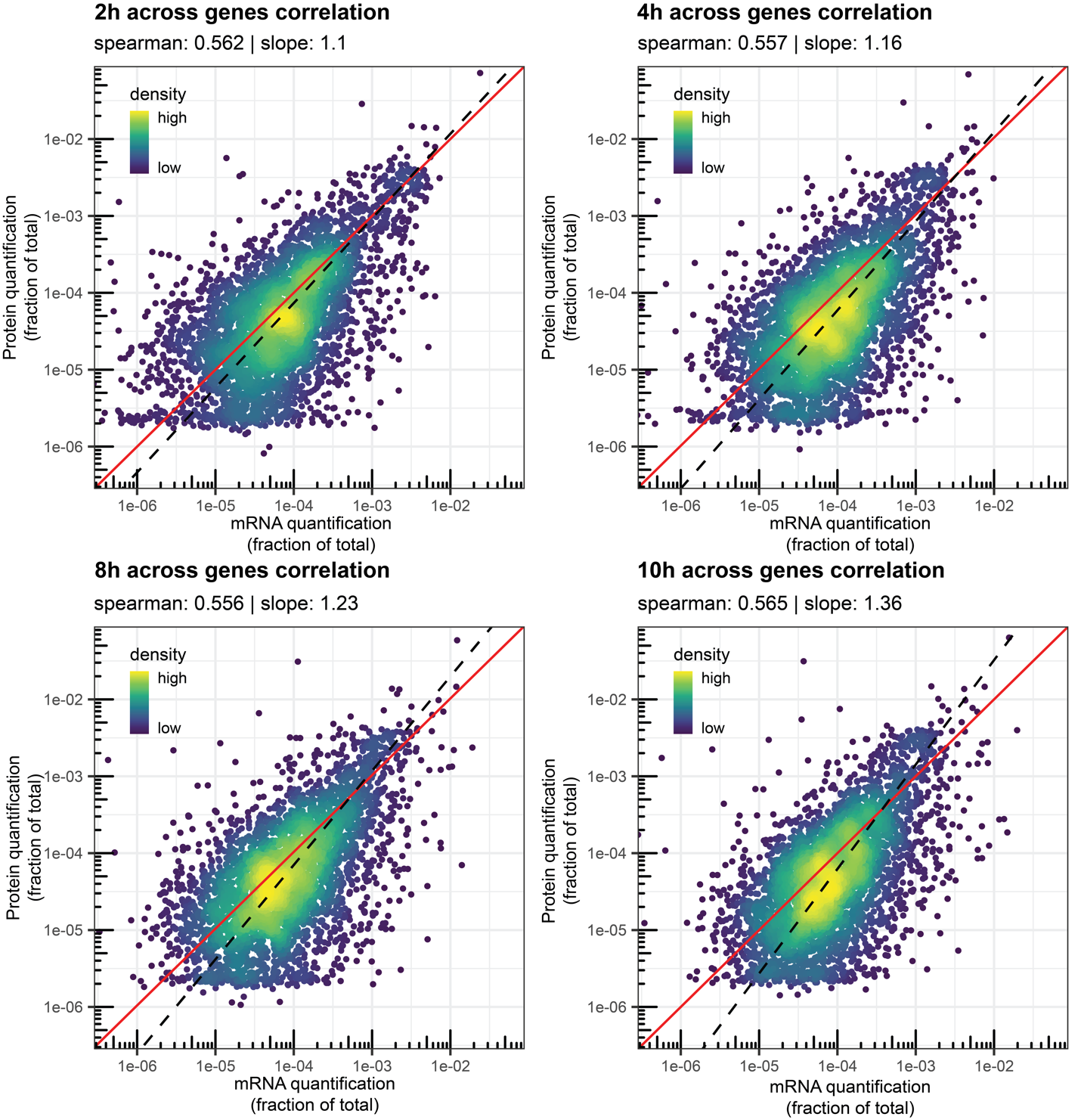

**Fig. S5 | Across genes correlation of mRNA and protein fractions at 2h, 4h, 8h, and 10h time points.** Correlation of the mean mRNA and protein levels across the indicated time point. Each dot represents the mean protein and mRNA expression from a single gene. The dashed black line indicates the linear regression of the data, with the slope indicated above the plot. The red line is the y=x diagonal.

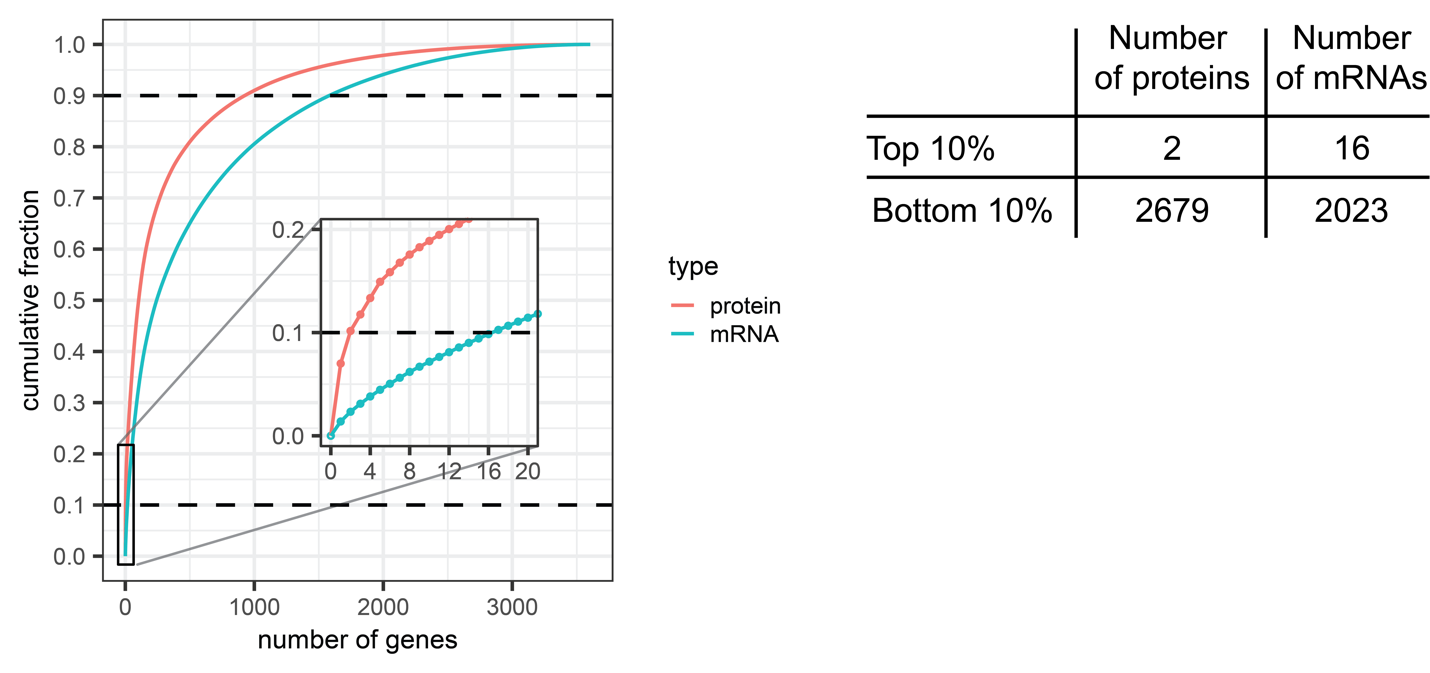

**Fig. S6 | Differences in dynamic range between transcriptomics and proteomics.** Cumulative fraction of proteomics and transcriptomics summing up to 1, for all genes quantified in both datasets. The dashed lines indicate the top and bottom 10% of expression, and the number of genes included in these cutoffs for the different datasets is included in the table.

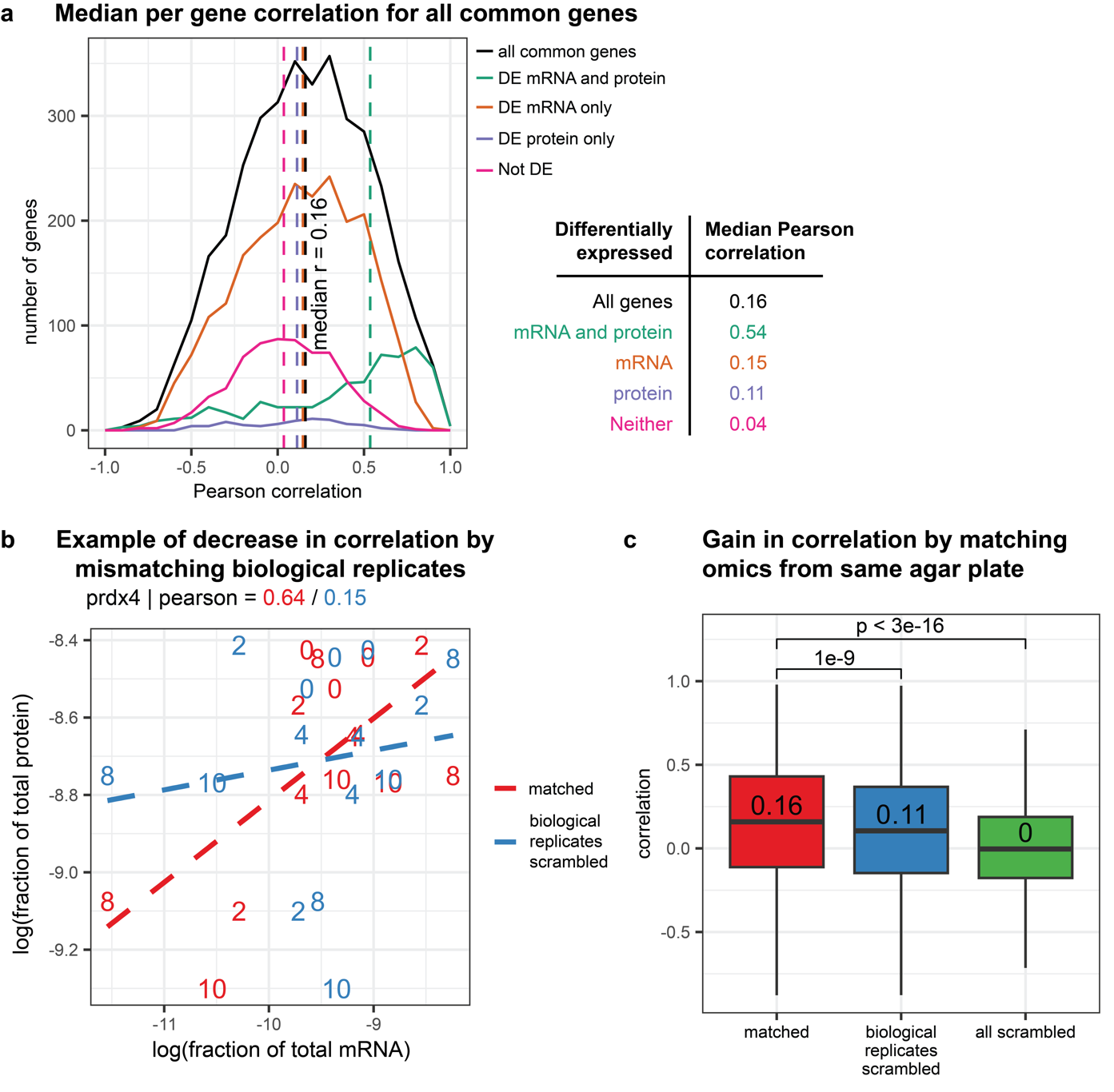

**Fig. S7 | Per gene mRNA and protein quantification. a** Distribution of per gene Pearson correlations for all common genes in the transcriptomics and proteomics datasets, and genes grouped by differentially expression in either of the datasets. The median Pearson correlation for each group of genes is visualized with a dashed line, and is noted in the table on the right. **b** Difference in Pearson correlation for prdx4 gene when mismatching biological replicates. Each replicate is plotted based on the mRNA abundance and protein abundance of either that same replicate (red) or protein abundance of another replicate from the same time point (blue). Linear regression indicated with dashed lines matching the color of the replicates, with a Pearson correlation indicated above the plot. **c** Boxplots of per gene correlation for all genes in the proteomics and transcriptomics datasets, either where the data is matched to the same biological replicate from the same plate (red), or where the data is matched to different biological replicates from the same time point (blue), or where the data is completely scrambled, i.e. protein data from one biological replicate and time point, matched to mRNA data from another biological replicate and time point (green). P-values from two-tailed t-test for indicated pairwise comparisons.

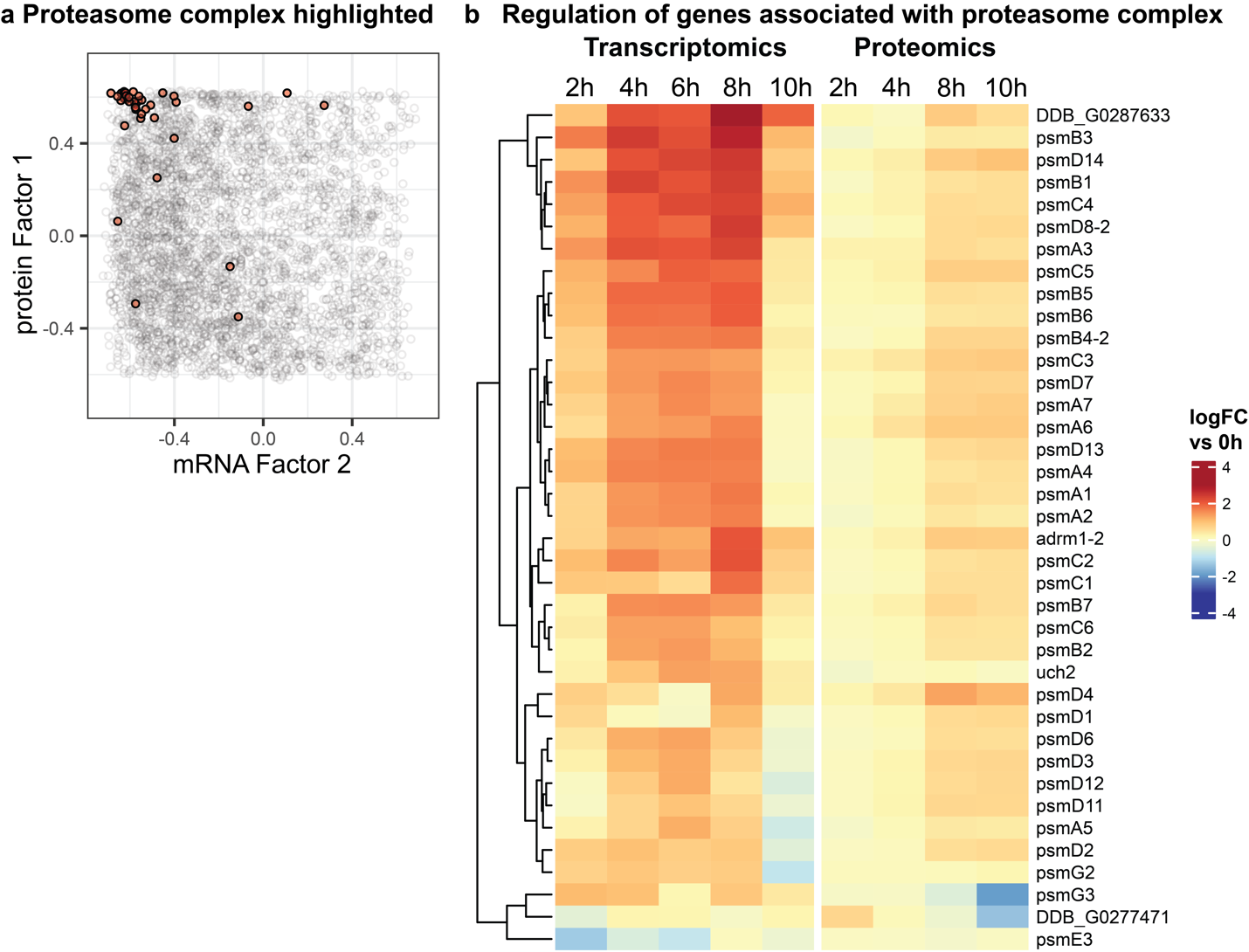

**Fig. S8 | Regulation of the proteasome complex during development. a** Highlight of members of the proteasome complex in mRNA factor 2 by protein factor 1 plot. In light-grey the distribution of all genes is shown. Genes in the top-left are associated with high Factor 1 at the protein modality, and low Factor 2 at the mRNA modality **b** Regulation of proteasome complex genes in transcriptomics and proteomics datasets in log fold change (logFC) versus the 0h time point.

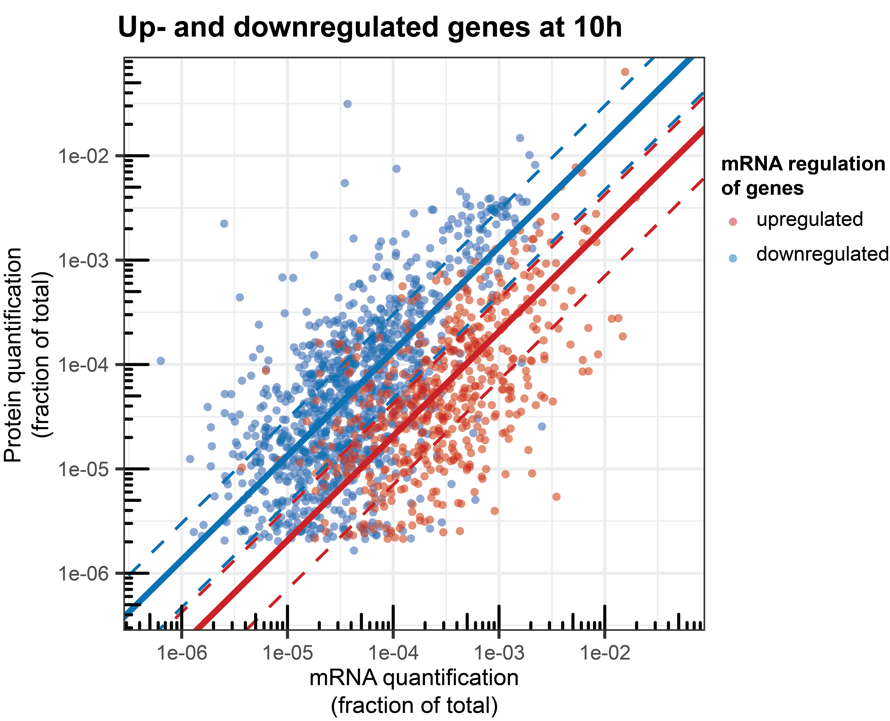

**Fig. S9 | Fraction of protein to mRNA for the 10h time point.** Across genes correlation of genes quantified in the transcriptomics and proteomics datasets. Each gene is plotted by the mean mRNA level and mean protein level at the 10h time point. Genes are included for which the mRNA is upregulated at the 10h time point (red) or genes for which the mRNA is downregulated at the 10h time point (blue). The median protein to mRNA ratio for each set of genes is indicated with the solid line in matching color. The 25th and 75th percentile are indicated with dashed lines.

3.	ProteomeXchange Dataset PXD023404 https://proteomecentral.proteomexchange.org/cgi/GetDataset?ID=PXD023404.
